# Gut Microbiome Prediction: From Current Human Evidence to Future Possibilities

**DOI:** 10.1101/2022.11.16.516694

**Authors:** Rinku Pramanick, Rajesh Kumar Gazara, Rafi Ahmad

## Abstract

The gut microbiome is an indispensable component of the human body. Alteration in the gut microbiota composition leads to various diseases such as obesity, Type 2 Diabetes, Inflammatory Bowel Syndrome, and depression. Microbiome-based precision tests offer a futuristic strategy for wellness and longevity. However, this approach is limited by the lack of definition of a healthy microbiome in different populations and accurate disease prediction.

In this study, we aimed to capture the healthy gut microbiome for different populations using the xNARA Gut Profile Test kit and in-house built proprietary algorithm and reference databases. We found that gut microbiome of different populations from India, UAE, and Singapore varied significantly, indicating a distinct geographic gut microbiome signature and Gut Health Index. The gut microbial diversity as measured by the Shannon index revealed UAE had significantly greater alpha diversity than India and Singapore. *Prevotella copri* (19.27%), *Faecalibacterium prausnitzii* (4.08%) and *Levilactobacillus brevis* (4.0%) were the predominant species in the Indian gut. *Faecalibacterium prausnitzii* (8.54%), *Blautia obeum* (8.10%), and *Phocaeicola vulgatus* (4.6%) were primarily present in Singapore participants whereas *Prevotella copri* (14.92%), *Blautia obeum* (6.09%) and *Roseburia intestinalis* (5.81%) were present in UAE participants. Beta diversity indicated the gut microbiota of Indian-origin participants in Singapore and UAE clustered with the indigenous inhabitants of Singapore and UAE. This highlights that geographic location has a profound effect on shaping the gut microbiome architecture than ethnicity. Regional diet and lifestyle could be crucial factors responsible for shaping the gut microbiome. The prediction accuracy of the xNARA Gut algorithm ranged from 66.66-100% when matched with the blood reports. Participants agreed with the xNARA disease risk outcomes for metabolic conditions (60%-100%), gastrofitness (62.5%-100%), mental health (50%-100%), skin conditions (50%-100%) and physical fitness (50%-100%). These observations imply the promising role of gut-based personalized diet and probiotic recommendations for lifestyle and wellness management.

## 1. Introduction

Communities of different microbes residing together in a particular habitat are called the microbiome. Such communities do exist in different sites of the human body like the respiratory tract, oral cavity and gastrointestinal tract (Costello et al, 2009). The gut microbiome is a community of microorganisms living in the gastrointestinal tract. The gut microbiota consists of approximately 500-1000 species of microbes. The ordinary human gut microbiota contains of main phyla, specifically Bacteroidetes and Firmicutes (Jandhyala et al, 2015).

The microbiota of the gut plays a significant role in regulating health and hence becomes an essential part of the study. Microbes help to break down complex carbohydrates, metabolize nutrients, and synthesize vitamins and compounds that enhance our immunity. The normal gut microbiome imparts specific characteristics in nutrient metabolism, drug metabolism, and immunomodulation and improves the integrity of gut mucosal barrier (Jandhyala et al, 2015). Certain gut bacteria play a significant role in weight management and exercise regimen. The diversity of gut bacteria can also affect the skin through the gut-skin axis.

The intestine microbiota is specially composed of beneficial and pathogenic bacteria (Sassone-Corsi & Raffatellu, 2015). Mostly, the microorganisms live in a symbiotic relationship in human microhabitats. The pathogens can cause illness when their number increases, therefore, a balanced composition of gut microbes is crucial for the prevention of diseases as well as physical and mental well-being (Bull & Plummer, 2014). Thus, the makeup of our gut microbes determines the risk of diseases such as IBD, diabetes, obesity, and cardiovascular diseases (Aschard et al, 2019). Through the gut-brain axis (the relationship between the gut and the nervous system); certain bacteria in the gut can affect mental health issues such as anxiety and depression (Simpson et al, 2021).

Every individual has their gut microbiome signature, depending on their geographic location and lifestyle. The geographic location seems to play one of the major roles leading to the variations in the gut microbiota. Several studies have proven that populations in higher latitudes tend to have larger body mass in comparison with populations in decreased latitudes. This is probably mediated via the means of gut microbial communities that modulate energy extraction and fat storage (Suzuki & Worobey, 2014). Moreover, the quality and quantity of the gut microbiome determine its function. Any deviation from the healthy gut microbiome composition, known as dysbiosis, is a profound indicator of disease onset and/or progression (Fan & Pederson, 2021). Dysbiosis effects by a peculiar ratio of commensal and pathogenic bacterial species (Belizário and Napolitano, 2015). The imbalance of gut microbes influences the fitness of the host in many ways, including energy absorption, choline, short-chain fatty acids, gut-brain axis, bile acids and consequently results to several adverse effects on health (Chen & Wang, 2021). With the rapid advances of next-generation sequencing and bioinformatic software, microbiome research has boomed in the last decade. This led to an enormous interest in deciphering the potential of the gut microbiome in preventive and therapeutic medicine and nutrition (Jain, 2020; Singh et al, 2017). Inter-individual deviations in the gut microbiome opened the scope for personalized microbiome tests. xNARA gut health kit is devised for personalized microbiome tests, giving a complete gut microbial profile along with risk assessment for 50+ health conditions encompassing physiology, mental health, aesthetics, and overall health.

Personalized microbiome testing requires defining a healthy reference microbiome for different geographic locations (He et al, 2018). In this study, we aimed to capture the healthy gut microbiome for different populations using 16s rRNA V3-V4 regions to identify population specific gut microbiome features.

## 2. Material & Methods

### 2.1.1 Study Participants

The study participants were recruited as per the following inclusion and exclusion criteria. Participants willing to give stool samples and participate in the study were recruited. The participant data was declassified and anonymized for personal data protection.

#### Inclusion criteria

➢ Apparently healthy and asymptomatic adults
➢ Age: 15-70 years
➢ BMI-18.2 to 25
➢ No complaints of any infection or disease

#### Exclusion criteria

➢ Pregnant ladies
➢ Nursing women
➢ Subjects with pacemakers and stents
➢ Subjects with therapeutic category of antineoplastics, steroidal drugs and antibiotics
➢ Used Antibiotics in the last 3 months
➢ Chronic disease (HIV, CKD, Cirrhosis, Acromegaly, active hyperthyroidism etc.)
➢ Cancer and anticancer treatment in the last 5 years
➢ Psychiatric disorders (bipolar disorder, schizophrenia, OCD)

### 2.2 Sample Collection

Participants were asked to collect the stool samples using the easy-to-use xNARA Gut Profile test kit (Fig. 1). Instructions to collect the sample were clearly printed with an easy-to-follow graphical presentation. Participant samples in DNA/RNA Shield (Make: Zymo Research) solutions were sent to xNARA testing labs.

**Fig. 1:**
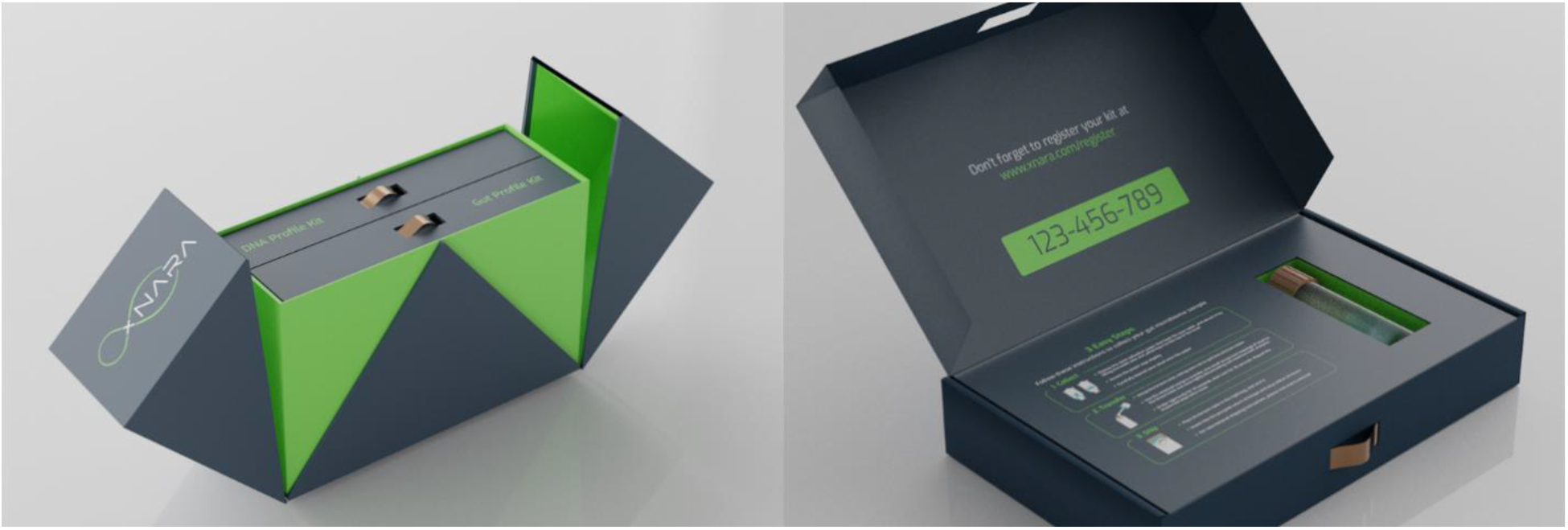
xNARA Gut Profile Test Kit.

### 2.3 Sample processing and DNA extraction

Microbial Genomic DNA was extracted according to the instruction on MO BIO Powersoil DNA extraction kit (Qiagen, Catalog No. 12888).

### 2.4 Amplicon Sequencing

Extracted DNA were analysed using next-generation sequencing (NGS) for assessment of the faecal microbial composition. The 16s rRNA V3-V4 libraries were sequenced on Illumina Mi-Seq using 2×300 bp chemistry yielding approximately 50,000 paired end reads per sample.

### 2.5 Data Analysis and Statistics

#### 2.5.1 Sequence data pre-processing

Sequencing reads quality was assessed using FastQC (v0.11.5) (https://www.bioinformatics.babraham.ac.uk/projects/fastqc/). Low bases and adapter sequences were removed by fastp (v0.23.2) (Chen et al, 2018) to obtain only high-quality and adapter free reads. The following parameters were used in fastp -t 10 -T 10 -u 40 along with minimum read length 50 bp and quality value for a base ≥ 20. FastQC package was run again and confirmed that no further filtering steps were required.

#### 2.5.2 Taxonomic Classification & Diversity Analysis

High quality and clean reads were processed with Kraken 2 (Wood et al, 2019) against the xNARA custom bacterial database for taxonomic assignment. The output of Kraken 2 was processed using Bracken 2 (Lu et al, 2017) to quantify relative abundances at species and genus taxonomic levels. The microbial Shannon diversity index (via diversity function) and Beta diversity were calculated using vegan package (Oksanen et al, 2013) at genus and species level in R. Beta diversity, Principal coordinates analysis (PCoA) derived from Bray–Curtis distances among samples was assessed by PERMANOVA using the adonis function of the R package vegan with 999 permutations. All the boxplots were plotted using ggplot2 package (Wickham et al, 2016) in R. Venn diagrams were plotted using Venn webtool (https://bioinformatics.psb.ugent.be/webtools/Venn/).

#### 2.5.3 Gut Health Index Score and disease-risk score

xNARA built region-wise (India, Southeast Asia and Middle East) models to calculate Gut Health Score. Data from healthy (n=2314) and unhealthy (n=283) study participants were used to establish reference values for these regions. Gupta et al, 2020 developed a Gut Microbiome Health Index (GMHI) (Gupta et al, 2020). xNARA did modifications in GMHI and constructed the Gut Health Score calculation pipeline, which can classify samples into poor, moderate and good for above mentioned regions separately. We calculated the Gut Health Index Score for all our samples using xNARA Gut Health Score pipeline.

xNARA developed an xNARA microbiome intelligent analytics workbench that contains a database of harmful and beneficial bacteria related to 59 conditions along with scoring a system-based disease risk scale. This xNARA workbench can classify metagenomics samples as poor, moderate, and good for various health conditions. We classified our samples using the xNARA microbiome intelligent analytics workbench. These health parameters encompass the four health-focused areas viz. Physiology, Mental Health, Aesthetics, and Overall Health. The xNARA disease risk outcomes were validated by taking feedback from the participants (n=50) and blood reports of a few volunteers (n=6).

## 3. RESULT

### 3.1 Characterization of participants recruited for the study

A total of 50 adult volunteers were enrolled in the study. The mean age of the participants was 35.89± 9.43 yrs. About 32% (16) of the participants were residents of India, 42% (21) from Singapore, and 30% (13) from UAE. Participant characteristics are elaborated in Table 1.

**Table 1:** Characteristics of participants recruited for the study

| Country | Characteristics | Mean | SD |
| --- | --- | --- | --- |
| India | Age (years) | 33.44 | 6 |
|  | Male (n= 9) | 32.73 | 6 |
|  | Female (n=7) | 32.35 | 6 |
|  | BMI | 24.78 | 3 |
| UAE | Age (years) | 35.85 | 6 |
|  | Male (n= 7) | 37.28 | 6 |
|  | Female (n= 6) | 35.85 | 6 |
|  | BMI | 27.53 | 7 |
| Singapore | Age(years) | 38.23 | 13 |
|  | Male (n= 9) | 38.24 | 13 |
|  | Female (n= 12) | 38.86 | 13 |
|  | BMI | 25.87 | 6 |

### 3.2 Microbial diversity varies across the countries

#### 3.2.1 Species richness

Species richness is the number of species present in a sample. The mean species richness of samples from the three countries were 219.7± 35.74 (India), 147.57±25.86 (Singapore), and 334.15±90.48 (UAE). Participants from UAE were characterized by a greater species richness in their gut whereas those from Singapore had the least. Species richness was substantially less in the Singapore population compared to the India and UAE samples.

#### 3.2.2 Alpha Diversity

Alpha diversity is the mean species diversity in a sample. Gut microbial diversity was measured by calculating the Shannon–Wiener diversity index of the samples from each population. The gut microbial diversity of the Singapore population was significantly lower than those of the UAE (p= 0.027). Likewise, the Shannon index differed significantly between populations from India and UAE (p= 0.05). However, there was no statistical difference between the Shannon index of the samples from India and Singapore (p= 0.85) (Fig. 2).

**Fig. 2:**
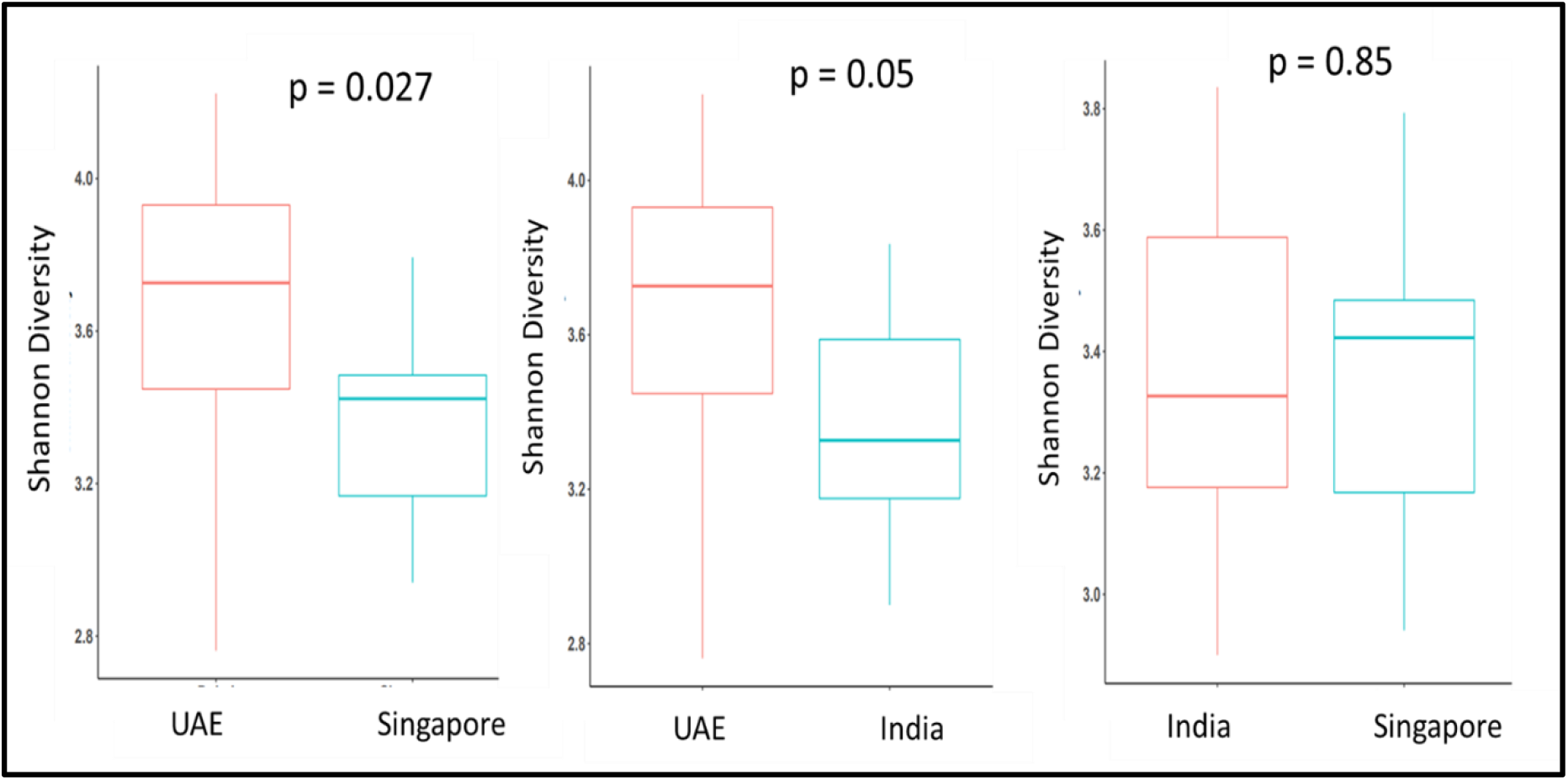
Alpha diversity of gut microbiome in different countries. The box plot demonstrates the Shannon alpha diversity index of UAE, India, and Singapore samples. The plot indicates that samples from Singapore are less diverse compared to the samples of participants from India and the UAE. Statistical significance was calculated using the Wilcox test. A P-value ≤ 0.05 was considered statistically significant.

#### 3.2.3 Beta Diversity

We noted statistically significant dissimilarities in the gut microbiota composition of populations from India, UAE, and Singapore (p= 0.001). The axis PCoA1 represents 24% variability and PCoA2 represents 14% variability constituting a combined variance of 38% between the samples (Fig. 3).

**Fig. 3:**
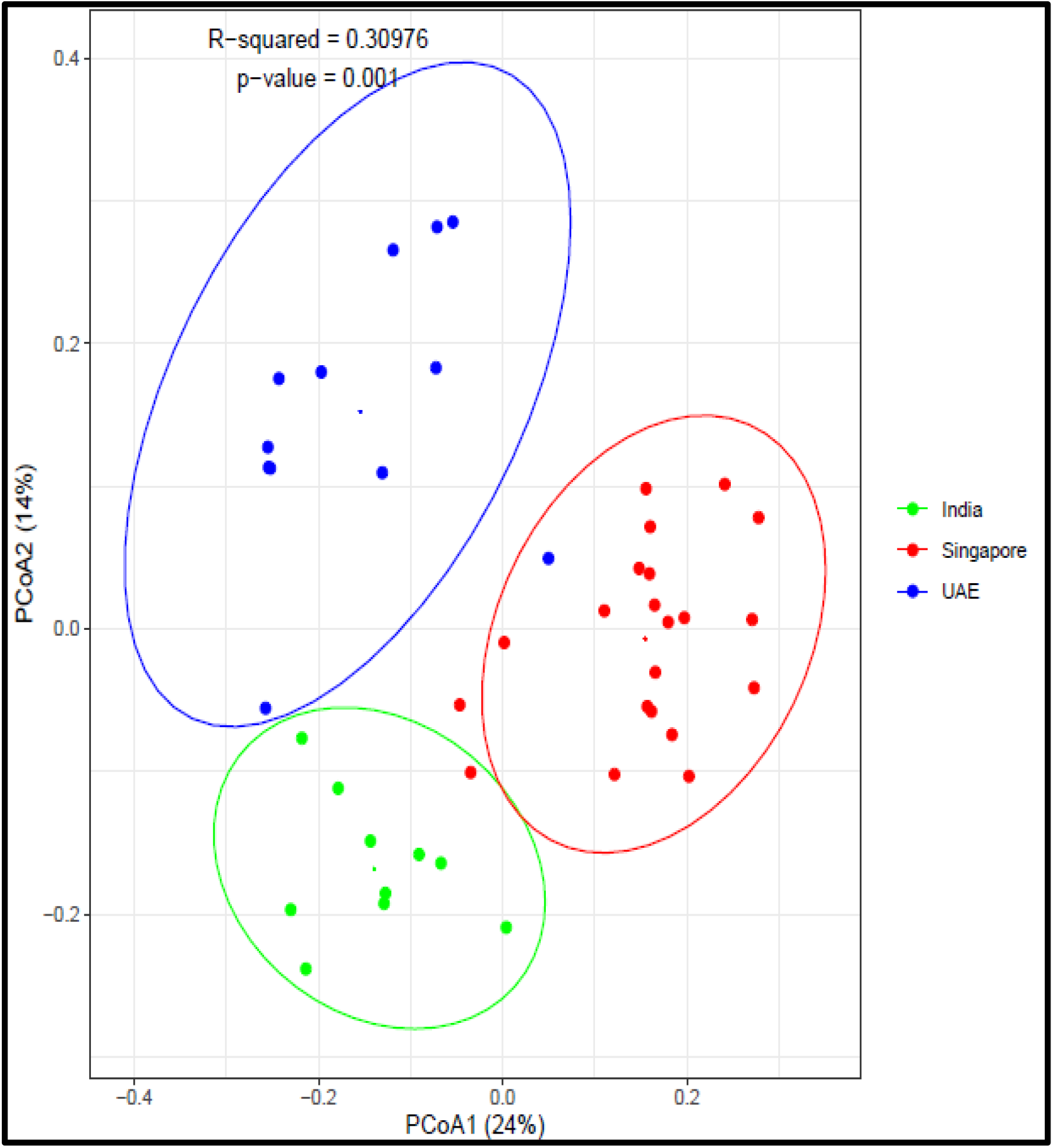
Beta Diversity. PCoA generated using Bray–Curtis distances indicates distinct clustering of samples from each population. Each point corresponds to an individual sample. For each population group, an ellipse around the centroid is depicted with a p value of 0.001.

#### 3.2.4 Geography Not Ethnicity is the Key Determinant of Gut Microbiome

Both geographic location and ethnicity can influence the gut microbial profile. In this study, few Indian-origin participants migrated to UAE and Singapore respectively. These immigrants stayed in these countries for at least a year. To investigate whether Indian-origin immigrants in UAE and Singapore clustered separately, we calculated the beta diversity of samples from different countries. Fig. 4a indicated the distribution of each Indian-origin immigrant sample with respect to the local inhabitants of UAE and Singapore. It is evident from Fig. 4b that the immigrants’ samples clustered along with the indigenous inhabitants of UAE and Singapore, respectively, leading to distinct segregation of population based on geographic location than ethnicity.

**Fig. 4:**
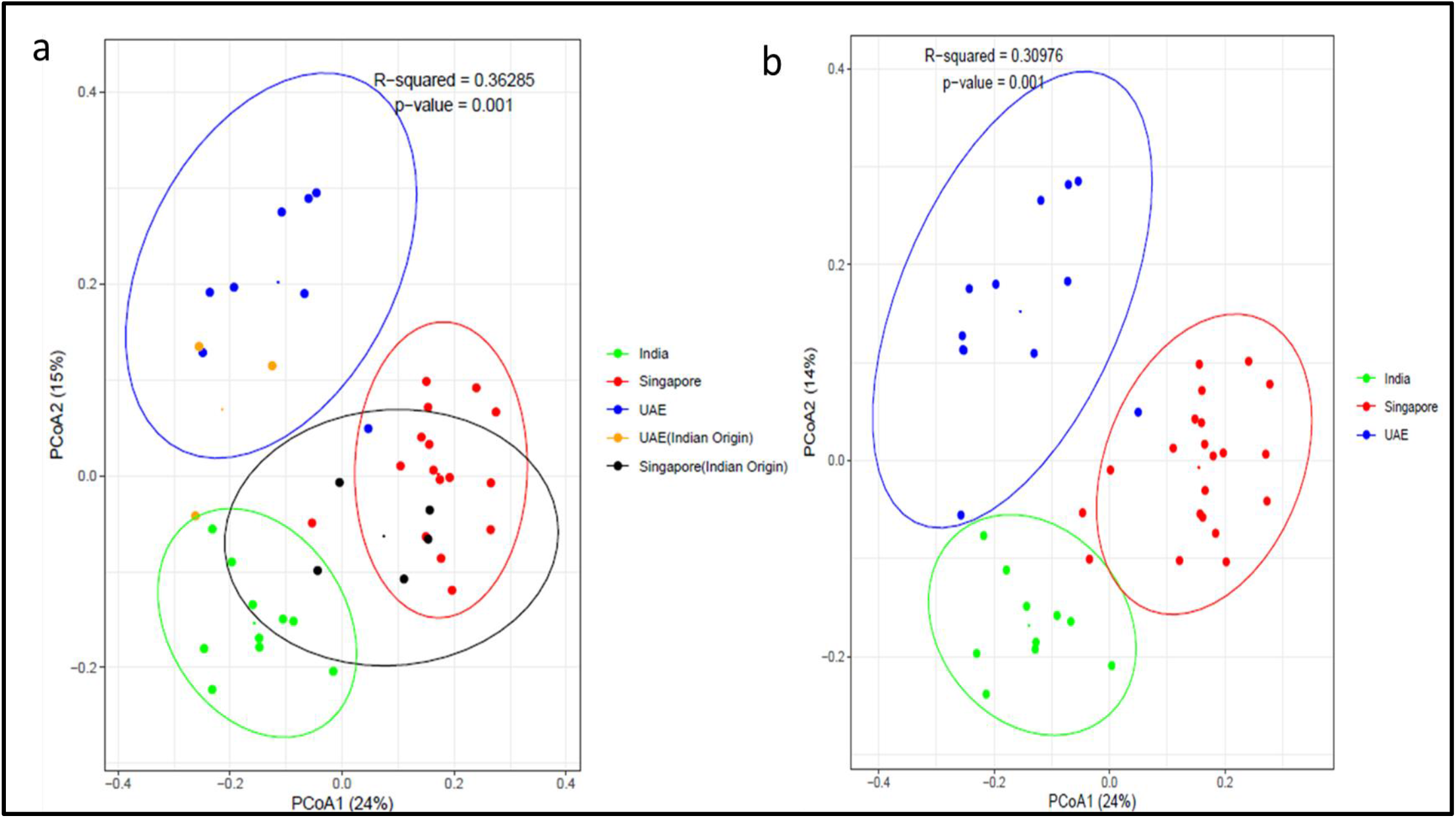
Beta diversity of gut microbiota of population from India, Singapore and UAE with immigrants. PCoA generated using Bray–Curtis distances. Each point corresponds to an individual sample. For each population group, an ellipse around the centroid is depicted. For each population group, an ellipse around the centroid is depicted with a p value of 0.001.

#### 3.2.5 Gut Microbiome Profile Across the Continent

Inhabitants of each region are characterized by a unique gut microbiome signature as shown in Fig. 5. *Prevotella copri* (19.27%), *Faecalibacterium prausnitzii* (4.08%) and *Levilactobacillus brevis* (4.0%) were the predominant species in Indian samples. People in Singapore predominantly harbor *Faecalibacterium prausnitzii* (8.54%), *Blautia obeum* (8.10%) and *Phocaeicola vulgatus* (4.6%) whereas *Prevotella copri* (14.92%), *Blautia obeum* (6.09%) and *Roseburia intestinalis* (5.81%) were present in UAE population.

**Fig. 5:**
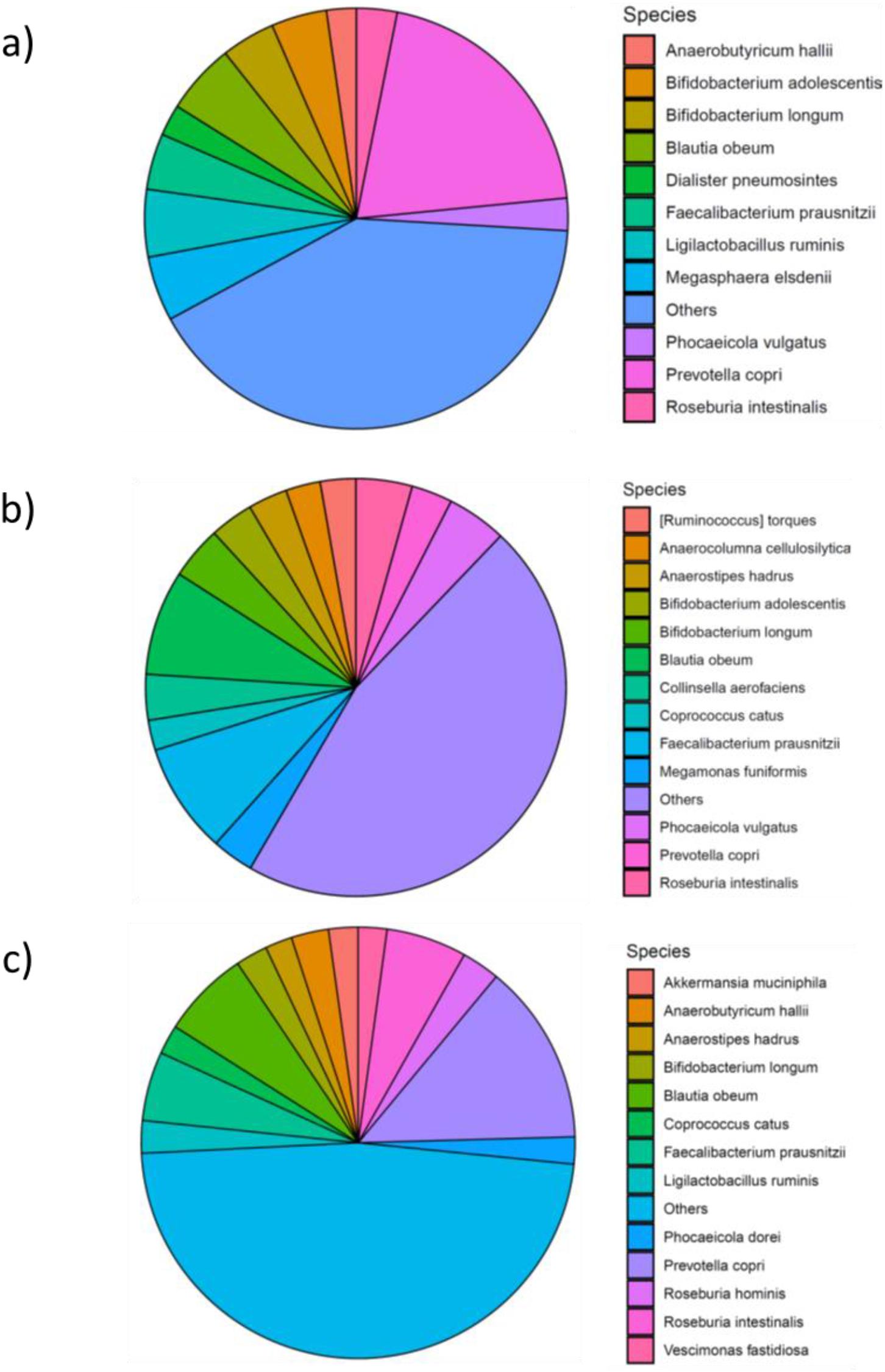
The gut microbiome profile of samples from (a) India, (b) Singapore (c) UAE. Species with a relative abundance of ≥ 2% are represented here.

#### 3.2.6 Differential Distribution of Gut Microbiome in The Continent

We observed the gut microbial profiles in different countries were characterized by the presence of unique genera and species (Fig.6a-d, Fig. S1). The Indian gut microbiome has *Dialister pneumosintes and Megasphaera elsdenii*, which are not present in other populations of this study. Likewise, *Anaerocolumna cellulosilytica, Collinsella aerofaciens, Ruminococcus torques*, and *Megamonas funiformis* were unique to Singapore and *Roseburia hominis, Akkermansia muciniphila, Vescimonas fastidiosa and Phocaeicola dorei* were present only in UAE population (Fig. 6b, Fig. S1).

**Fig. 6:**
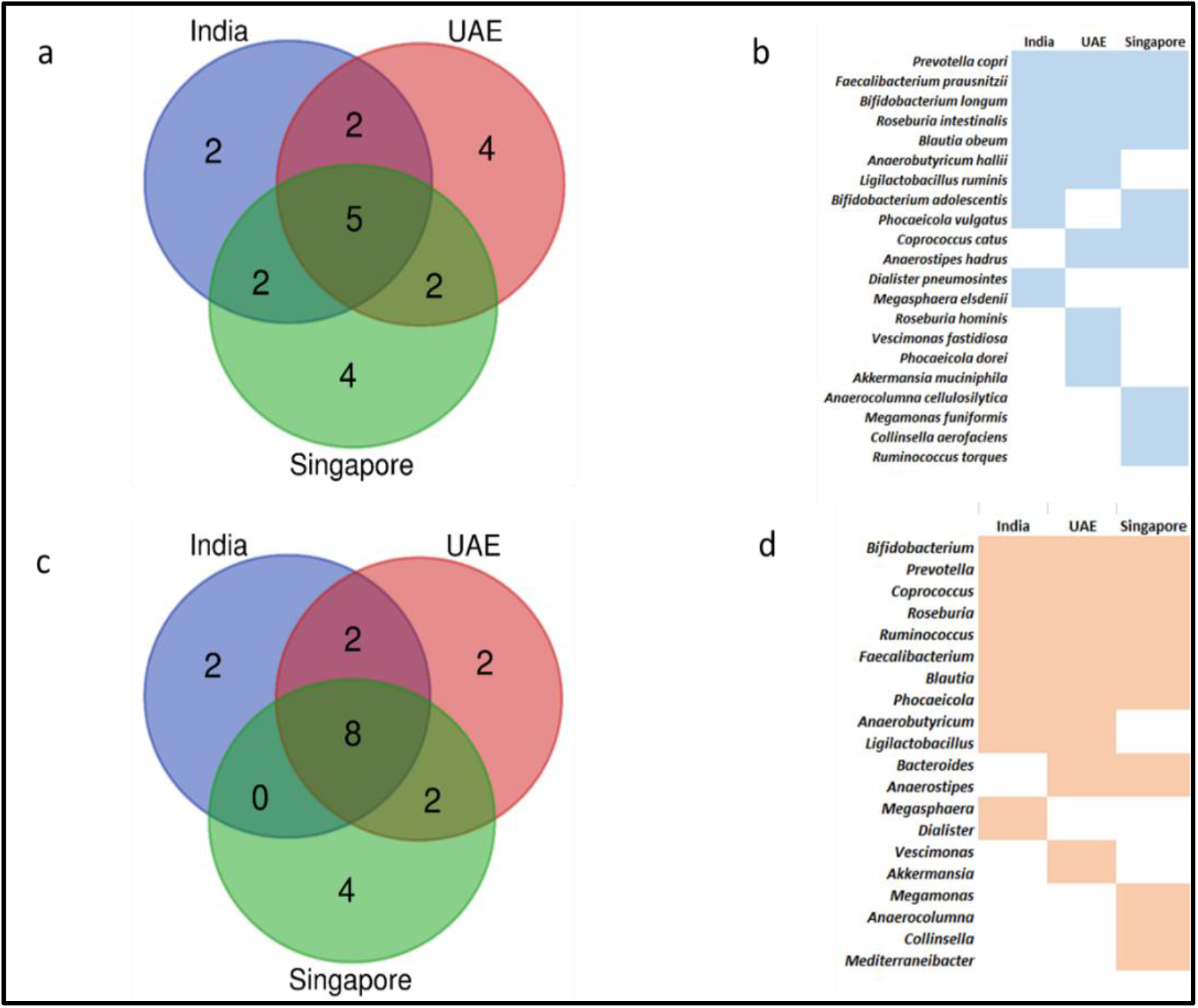
Common and unique gut microbiota present across the different countries. a) and b) species level and c) and d) genus level.

Apart from the unique species, we found five species were present across these countries. These species were *Prevotella copri, Blautia obeum, Roseburia intestinalis, Bifidobacterium longum, Faecalibacterium prausnitzii* (Fig. 6a&b). At the genus level, *Megasphaera* and *Dialister* were exclusively present in Indian gut microbiota, whereas *Megamonas, Anaerocolumna, Collinsella*, and *Mediterraneibacter* colonized the Singapore gut and UAE was characterized by the presence of *Akkermansia* and *Vescimonas* (Fig. 6c&d).

#### 3.2.7 xNARA Gut Health Index

The xNARA Gut Health Index indicates gut health status based on health-associated and unhealthy-associated bacteria. This index distinguishes the healthy samples from the unhealthy samples.

In the present study, we observed that the Gut Health Index varies for each geography like the gut microbiome profile. India’s mean Gut Health Index was 53.81±6.28, Singapore 40.71±18.32 and UAE 50.84±12.61which varied significantly among the groups (Fig. S2).

#### 3.2.8 xNARA Gut Health Test Disease Risk Prediction when compared with Blood report

The xNARA Gut disease risk algorithm evaluates the gut status for developing a particular disease and/or condition as good, moderate and poor. The xNARA Gut disease risk outcomes were matched with the blood results of a few study participants (n=6) in India (Table 2).

**Table 2:** Prediction Accuracy (%) of xNARA Gut Health Disease Risk prediction with blood results.

| Health Parameter | Prediction Accuracy (%) |
| --- | --- |
| Alcoholic liver disease | 83.33 |
| Vitamin B7 | 100 |
| Blood Lipid levels-HDL | 66.66 |
| Blood Lipid levels-TGL | 66.66 |
| Calcium | 83.33 |
| CVD | 66.66 |
| Kidney Stones | 83.33 |
| Prediabetes | 83.33 |
| Type II Diabetes | 100 |
| Vitamin A | 100 |
| Vitamin B1 | 100 |
| Vitamin B2 | 100 |
| Vitamin B5 | 100 |
| Vitamin B6 | 100 |
| Vitamin B9 | 66.66 |
| Vitamin D | 83.33 |
| Average | 86.45 |

#### 3.2.9 User Validation of the xNARA Gut Disease Risk Prediction

##### Predicting Major Metabolic Diseases

Type 2 Diabetes is associated with low-grade inflammation and microbiota present in the gut which significantly contributes to the pathogenesis of this disease. Timely diagnosis or prediction could offer better management of the disease. Based on the feedback, 88.88-100% of participants agreed with the outcome of the xNARA microbiome disease risk prediction (Table 3).

**Table 3:**
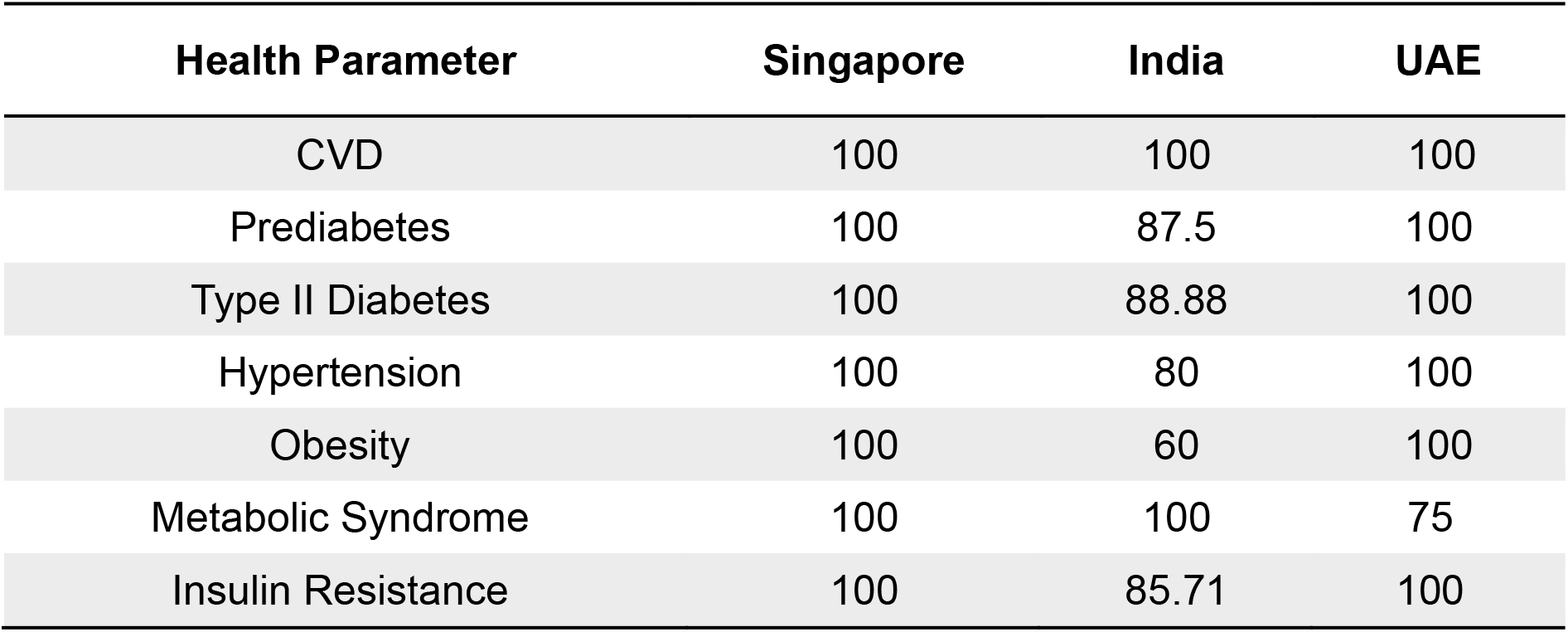
Prediction accuracy (%) of the xNARA gut microbiome test for metabolic conditions based on participant’ feedback

##### Predicting Gastrointestinal Disorders

An altered gut microbiota is an indicator of a malfunctioned gastrointestinal tract resulting in numerous anomalies such as diarrhea, bloating, IBD, constipation (Jalanka-Tuovinen et al, 2011; Carroll et al, 2012). The participant agreement for gastrointestinal disorders ranged from 62.5%-100% (Table 4).

**Table 4:** Prediction accuracy (%) of the xNARA gut microbiome test for gastrointestinal disorders based on participant’ feedback

| Health Parameter | Singapore | India | UAE |
| --- | --- | --- | --- |
| IBD | 100 | 90 | 100 |
| IBS | 66.67 | 100 | 100 |
| Crohn's Disease | 80 | 100 | 100 |
| Ulcerative Colitis | 80 | 88.88 | 100 |
| Constipation | 62.5 | 100 | 100 |
| Bloating | 66.66 | 100 | 75 |
| Flatulence | 83.3 | 88.88 | 100 |
| Diarrhea | 85.71 | 100 | 100 |

##### Predicting Mental Health Conditions

Clinical studies have shown that the gut microbiota of depressed patients is significantly different from that of healthy controls. Likewise, the gut microbiota is hampered during mental health conditions such as anxiety, post-traumatic stress disorder (PTSD) and migraine (Hemmings et al, 2017; Jiang et al, 2018). Maximum participants agreed with the results for depression across the countries (87.5%-100%) followed by migraine, anxiety, and PTSD (Table 5).

**Table 5:** Prediction accuracy (%) of the xNARA gut microbiome test for mental health conditions based on participant’ feedback.

| Health Parameter | Singapore | India | UAE |
| --- | --- | --- | --- |
| Depression | 87.5 | 90 | 100 |
| Anxiety | 57.14 | 80 | 100 |
| PTSD | 50 | 62.5 | 100 |
| Migraine | 57.14 | 87.7 | 100 |

##### Predicting Skin Conditions

The gut microbiota diversity could influence the skin through gut-skin axis. Participant agreements were highest for atopic dermatitis followed by vitiligo, psoriasis and acne results (Table 6).

**Table 6:** Prediction accuracy (%) of the xNARA gut microbiome test for skin conditions based on participant’ feedback

| Health Parameter | Singapore | India | UAE |
| --- | --- | --- | --- |
| Acne | 50 | 100 | 100 |
| Psoriasis | 100 | 75 | 100 |
| Atopic dermatitis | 100 | 100 | 100 |
| Vitiligo | 80 | 100 | 100 |

##### Predicting Physical Fitness Performance

The gut microbiota can play a role in the maintenance of a healthy body, as it can affect muscle strength through the gut-muscle axis. Based on the feedback, 50-100% of participants agreed with the outcome of the test (Table 7).

**Table 7:** Prediction accuracy (%) of the xNARA gut microbiome test for physical fitness performance based on participant’ feedback

| Health Parameter | Singapore | India | UAE |
| --- | --- | --- | --- |
| Active lifestyle | 66.67 | 90 | 75 |
| Aerobic Performance | 50 | 100 | 75 |
| Endurance | 100 | 100 | 100 |
| Muscle strength | 83.3 | 85.7 | 100 |

##### Predicting Sleep Quality

Gut microbiota can greatly impact sleep quality. The sleep prediction accuracy of xNARA Gut Health test for Singapore was 83.33% and 100% for India and UAE.

##### Predicting Food Intolerance

Gut microbiota affects the way our body reacts to gluten. The gluten intolerance prediction accuracy of the test was 100% for Singapore, India, and UAE. Lack of lactose metabolizing gut bacteria can lead to lactose intolerance. The prediction accuracy for lactose intolerance was 100% for Singapore and UAE and 90% for India.

## 4. DISCUSSION

The gut microbiota is considered as an additional human organ due to its indispensable role in determining our health (O’Hara AM and Shanahan, 2006; Baquero and Nombela, 2012). The gut microbial architecture determines the risk of diseases such as IBD, diabetes, obesity, cardiovascular diseases, brain health, sleep quality, skin health (Loubinoux et al, 2002; Mohajeri, 2018; Ley et al, 2006; Aron-Wisnewsky & Clément, 2016; Anderson et al, 2017; De Pessemier et al, 2021). Health and wellness tests based on gut microbiome are often limited by suboptimal prediction accuracy and ambiguous reference microbiome. One of the primary limitations of gut health tests is characterizing the healthy human microbiota.

In this study, we explored the gut microbiome of participants from India, Singapore, and UAE to identify gut microbiome features that are associated with these populations. We noted distinct gut microbiome composition for India, Singapore and UAE. The presence of *Dialister pneumosintes and Megasphaera elsdenii* were unique to the Indian gut, whereas *Anaerocolumna cellulosilytica, Collinsella aerofaciens, Ruminococcus torques*, and *Megamonas funiformis* were exclusively present in Singapore gut and *Roseburia hominis, Akkermansia muciniphila, Vescimonas fastidiosa and Phocaeicola dorei* were present only in UAE samples. Furthermore, the gut microbiota of samples from the same country clustered together irrespective of their ethnicity, suggesting a prominent influence of regional lifestyle over genetics. The local diet, environment and lifestyle play a vital role in framing the gut microbiota ecosystem (Leeming et al, 2019; Tasnim et al, 2017; Conlon & Bird, 2014). Such factors limit the applicability of gut microbiome for precision nutrition (Nogal et al, 2021). These findings mandate the need for region-specific baseline healthy gut microbiome reference. Using the respective baseline healthy reference from where the customer belongs will enable accurate reporting and precise microbiome prediction. Furthermore, accurate identification of the pathogens and the probiotic bacteria in the sample is crucial for robust results and appropriate health recommendations. To characterize and accurately predict the gut microbiome status xNARA has established regional healthy gut microbiome references for India, Southeast Asia, and the Middle East. Gut microbiome data of healthy samples and unhealthy samples were used to establish reference values for these regions. Likewise, xNARA has calculated the proprietary xNARA Gut Health Index (GHI) from healthy and unhealthy samples from these countries. xNARA GHI can predict gut health status and classify it as good, moderate, and poor. Each country has a unique xNARA Gut Health Index (GHI) as observed from this study.

xNARA microbiome intelligent analytics workbench evaluates the gut microbiome for 59 health conditions. The prediction accuracies for most of the diseases were outstanding when compared with blood reports. Similarly, participants from all three countries agreed with the gut test results. The dysbiotic microbiome in individuals could be restored through diet and lifestyle recommendations (Petrosino, 2018). Thus, it is promising to unravel the interplay of diet on the gut microbiome architecture of these participants before offering any personalized diet and lifestyle recommendations.

Using carefully selected beneficial probiotic strains confers potential health benefits against microbiome-associated diseases (Kim et al, 2019). However, the ‘one size fits all’ approach gave rise to the use of over-the-counter probiotics with little or no clinical benefits (Suez et al, 2019). One of the reasons for the inefficacy of such probiotics is due to inter-individual variations of the gut microbiota (Lozupone et al, 2012; Kim & Jazwinski, 2018). Every individual has a unique gut microbiome signature that changes over the period based on environmental and lifestyle changes (Kim & Jazwinski, 2018). A tailored made personalized probiotics formulation based on the current gut microbiome composition represents a promising approach for rehabilitating the gut microbiota against various health complications. With this regard, the xNARA Microbiome based gut health evaluation will offer potential preventive strategies by rebalancing the gut microbiota through diet, lifestyle and clinically proven probiotics precisely personalized for every individual.

## Author contributions

RA conceived the study and designed the analysis. RP and RKG performed the analysis. RP and RKG wrote the first draft and RA reviewed the manuscript. All authors contributed to the article and approved the submitted version.

## Funding

This work was funded by xNARA BioLogics Pte. Ltd, Singapore.

## Conflict of interest

The authors report no conflict of interest.

## Acknowledgments

The authors wish to thank Dr. Sukithar Rajan for his bioinformatics support in the initial phases of the study. The authors would also like to thank all the trial participants who trusted in the study so that the advantages of the results could reach the community on a large scale.

## Notes

### Competing Interest Statement

The authors have declared no competing interest.

