## supplementary file for "Gut Microbiome Prediction: From Current Human Evidence to Future Possibilities"

**Supplementary Figures**

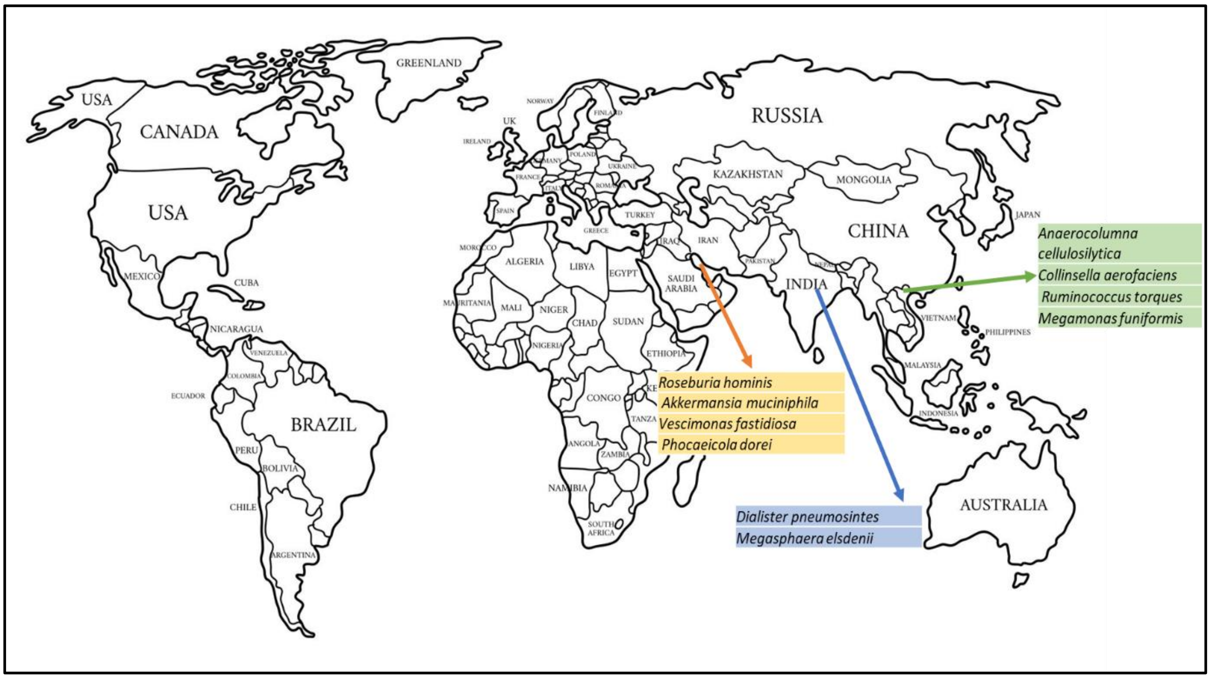


Fig.S1: Unique gut microbial species in diverse populations around the world.

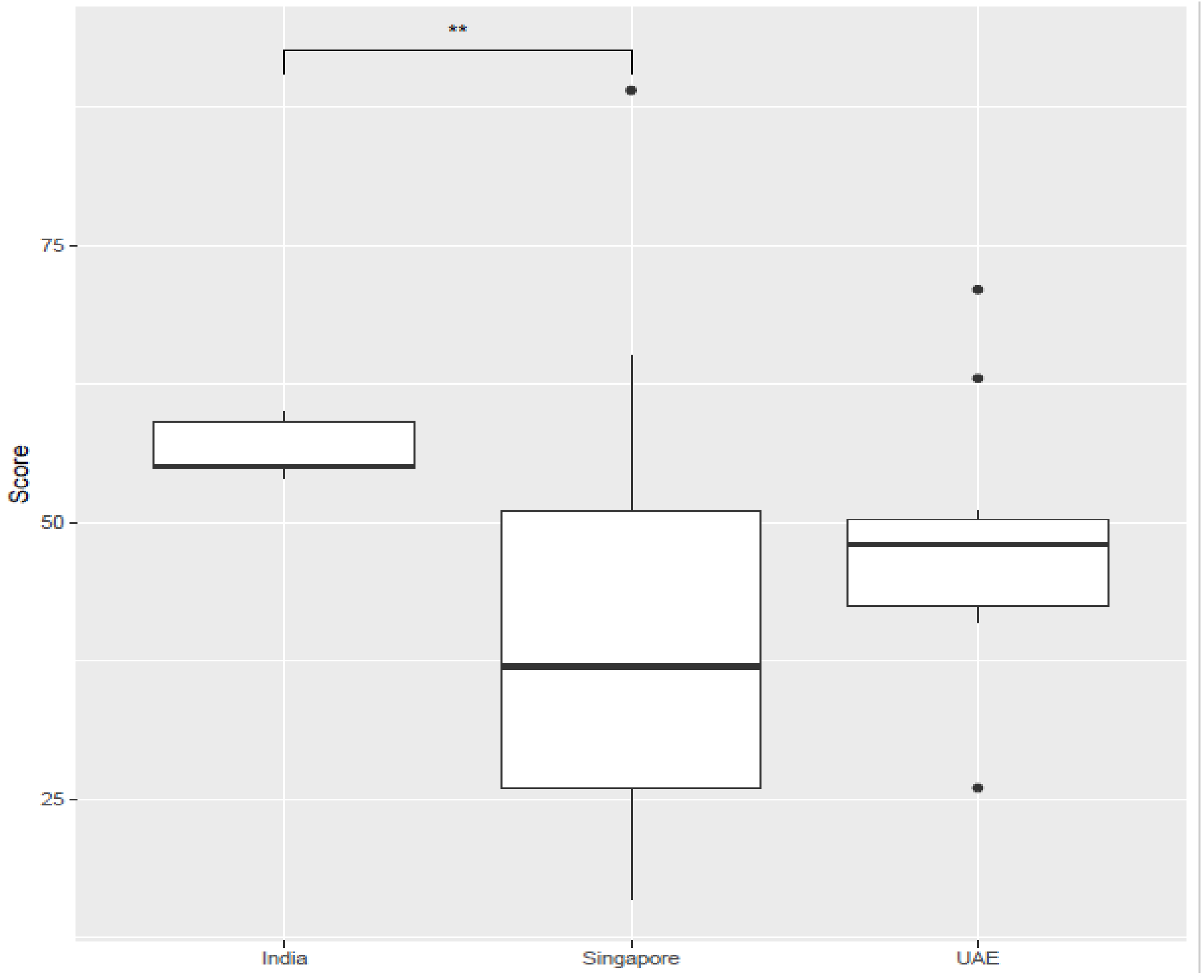


Fig.S2: Gut Health Index of the population of India, Singapore, and UAE
